# Evaluation of Silica and Bioglass Nanomaterials in Pulp-Like Living Materials

**DOI:** 10.1101/2024.10.07.616939

**Authors:** Daline Mbitta Akoa, Anthony Avril, Christophe Hélary, Anne Poliard, Thibaud Coradin

## Abstract

Although silicon is a widespread constituent in dental materials, its possible influence on teeth formation and repair remains largely unexplored. Here we have studied the effect of two silicic acid-releasing nanomaterials, silica and bioglass, on a living model of pulp consisting of dental pulp stem cells seeded in dense type I collagen hydrogels. Silica nanoparticles and released silicic acid had little effect on cell viability and mineralization efficiency but impacted metabolic activity, delayed matrix remodeling and led to heterogeneous cell distribution. Bioglass improved cell metabolic activity and led to a homogenous dispersion of cells and mineral deposits within the scaffold. These results suggest that the presence of calcium ions in bioglass is not only favorable to cell proliferation but can also counter-balance the negative effects of silica and silicic acid. Both chemical and biological processes should therefore be considered when investigating the effect of silicon-containing materials on dental tissues.

## INTRODUCTION

In the field of dentistry, biomimetic tissue engineering strategies based on the design of cellularized or living materials recapitulating key features of a targeted tissue has mainly focused on the dentin-pulp complex.^1–3^ Among the three main tissues of the dental organ, enamel, the outer tissue of the tooth, is not cellularized because ameloblasts, that are responsible for its formation, die at the stage of tooth eruption.^4^ Since ameloblast-like cell lines are difficult to obtain, biomimetics approaches of enamel reconstruction have so far focused on acellular models.^5^ In contrast, the most inner dental tissue, pulp, is an innervated and vascularized, non-mineralized tissue, hosting a wide variety of cells, including fibroblasts, immune cells as well as neural crest-derived stem cells.^6^ Dentin, a porous tissue with a composition close to bone, lies between enamel and pulp.^7^ The “primary” dentin is built-up by odontoblasts during early tooth development. Thereafter, odontoblasts localized at the dentin-pulp interface synthesize the “secondary” dentin all along the tooth life. The “tertiary” dentin is formed in response to tooth lesions. In case of caries reaching the surface of dentin only, the remaining odontoblasts are activated and form a “reactionary” dentin. When the caries reach the pulp and odontoblasts are thus destroyed, dental pulp stem cells (DPSCs) are recruited to differentiate to odontoblast-like cells able to promote the formation of a “reparative” dentin.^8^ DPSCs are therefore the main focus of tissue engineering strategies for dentin repair.^9–11^ Besides their direct therapeutic applications, DPSC-laden scaffolds can be used as *in vitro* tissue models to better understand more fundamental processes or drug/safety evaluation platforms.^12,13^

On this basis, several pulp-like 3D models consisting of DPSCs encapsulated within hydrogels made from natural or synthetic polymers have been described.^14–16^ To recreate the dentin-pulp-interface, these hydrogels are often placed within a dentin slice.^17^ To increase the physiological relevance of these models, co-cultures of DPSCs with other relevant cell types such as endothelial cells have also been reported.^18^ The pulp extracellular matrix being predominantly composed of type I and type III collagen,^6^ type I collagen-based hydrogels appear particularly suitable to design biomimetic hosts for DPSCs.^19–21^ However, although pulp is considered as highly hydrated, the corresponding collagen concentration (25 w:w %, *i.e*. 250 mg.mL^-1^)^6^ is much higher than those traditionally used to design cell-laden collagen hydrogels (2-10 mg.mL^-1^).^22^ In this context, the plastic compression process, that allows to prepare cellularized dense collagen hydrogels,^23^ has appeared as a method of choice to prepare pulp-like living materials.^24,25^

Silicon-based materials are widely used in dental repair,^26–28^ including cements^29^ and micro/nanoparticles used as fillers in composite resins.^30^ These materials are likely to release silicon species during material deposition, ageing or degradation on dental cells. In particular, calcium silicates are used as pulp capping materials, *i.e*. applied on injured pulp to attenuate inflammation and promote DPSCs differentiation into odontoblast-like cells, ultimately favoring remineralization.^31–32^ These materials are able to release Ca^2+^, OH^-^ and Si(OH)4 (silicic acid) species, similarly to bioglasses widely used in bone regeneration.^33,34^ However, whereas the possible role of silicic acid on bone health and repair has been extensively studied,^35,36^ the potential influence of silicic acid on dental repair and, more specifically, on DPSCs differentiation and mineralization ability, has been very sparingly studied.^37–39^ In particular, our previous research evaluated the effect of silicic acid at physiological (10 μM) and supraphysiological (100 μM) concentrations in pulp-like 3D models consisting of DPSCs seeded in dense collagen hydrogels prepared by the plastic compression method.^39^ This study suggested that Si(OH)4 had no direct effect on DPSCs but could instead interact with the collagen matrix and modify cell-matrix interactions, resulting in impaired cell migration and mineralization inhibition at high concentration.

However, in a clinical situation, Si(OH)4 is not present at a constant dose but is progressively released from materials. Moreover, in the case of bioglass or calcium silicates, Ca^2+^ ions are released concomitantly. On this basis, the aim of this work was to evaluate how the kinetics of silicic acid release and the presence of calcium ions could influence the behaviour of DPSCs embedded into dense collagen hydrogels. For this purpose, silica and bioglass nanoparticles were incorporated into the pulp-like living materials. The viability, metabolic activity, differentiation, organization and mineralization ability of the DPSCs were assessed as a function of particle composition and concentration. Results were discussed taking into account both the particulate and dissolved forms of silica and bioglass and considering concomitant biological and chemical processes.

## MATERIALS AND METHODS

### Silica Nanoparticles Preparation

Silica nanoparticles (Si) were synthesized using the Stöber process.^40^ In the standard synthesis protocol, 0.5 mL of tetraethylorthosilicate (TEOS 98%, Sigma-Aldrich) was rapidly mixed with 2 mL ethanol. Next, this TEOS/ethanol mixture was introduced into a solution containing 100 mL ethanol and 20 mL NH4OH (30%, Merck), and stirred for 2 h at room temperature. The resulting nanoparticles were collected by centrifugation, dispersed by ultrasonication and alternately centrifuged three times with ethanol and deionized water to remove any unreacted precursors. The resulting white solid was freeze-dried for 24 hours.

### Bioglass Nanoparticles Preparation

The synthesis of bioactive glass nanoparticles (BG) was conducted via a previously established methodology.^41^ Initially, two distinct solutions were prepared at ambient temperature. Solution 1 was formulated by swiftly adding 2.34 mL of TEOS to 20 mL of absolute ethanol, while solution 2 was composed of 11.7 mL of deionized water, 17.5 mL of absolute ethanol, and 1.7 mL of ammonium hydroxide. Following 30 minutes of mechanical stirring, solution 1 was introduced into solution 2 and allowed to react for 2 hours. Subsequently, 5 g of calcium nitrate, ((Ca(NO3)2).4H2O) was incorporated into the resulting turbid mixture and subjected to stirring for 22 hours. The resulting white suspension was centrifuged and subjected to sequential washing steps involving ethanol and water to eliminate residual reagents. The resulting white solid was dried overnight at 60°C, followed by a heat treatment at 650°C for 3 hours.

### Nanoparticles characterization

Nanoparticle size and morphology were studied by Dynamic Light Scattering (DLS) on a Zetasizer nano-ZS90 and scanning electron microscopy (SEM) using a Hitachi S-3400 N microscope operating at 60 kV. Chemical composition was studied by Attenuated Total Reflectance – Fourier Transform InfraRed spectroscopy (ATR-FTIR) using a PerkinElmer Spectrum 100 equipment. Wavelength-dispersive X-ray fluorescence (WDXRF) measurements were performed on a Bruker S8 Tiger apparatus to determine the % of silicon and calcium atoms in the BGs.

### Human Dental Pulp Stem Cells

Teeth were extracted from patients aged 15-20 years for orthodontic reasons at the AH-HP Bretonneau, according to the ethical guidelines established by French bioethics law (IRB agreement 00006477 and n° DC-2009-927, Cellule Bioéthique DGRI/A5), under an opt-out consent model. Human Dental Pulp Stem Cells (hDPSCs) were recovered and expanded following a previously reported protocol.^42^ Cells were used at passage 4.

### Dense Collagen Hydrogel Preparation

Solutions of Type I collagen (4 mg.mL^-1^) extracted from young rat tails in a 20 mM acetic acid solution was mixed with 5X DMEM (Dulbecco’s Modified Eagle Medium) and adjusted to the physiological pH with 0.1 M NaOH. DMEM media supplemented with 10% fetal bovine serum and 1% penicillin/streptomycin containing either silica or bioglass nanoparticles, without or with suspended hDPSCs, were added to the collagen solution. Final collagen concentration was 1.6 mg.mL^-1^, final silicon concentrations were 100 μM or 1000 μm for both types of particles and the pre-compression concentration of hDPSCs was 2 × 10^6^ cells.mL^-1^.^25^ The mixture was placed into a 24-well plate and incubated at 37°C under 5% CO2 for 30 min. Recovered gels were deposited on a stack of blotting paper, nylon and stainless-steel mesh and loaded with a compressive stress of 1 kPa for 5 min, yielding dense collagen gel (DC).^24^

### Cell culture experiments

The hDPSCs encapsulated in plastically compressed dense collagen hydrogels were grown in a mineralizing induction medium (MIM) made of 300 µM L-ascorbic acid sodium salt, 10 nM dexamethasone, 10 mM β-glycerophosphate, 10% fetal bovine serum, and 1% penicillin/streptomycin. Cell-laden collagen gels were cultured in 6-well plates using 5 mL of culture medium per well. Gels were cultured for 28 days, with differentiation media being renewed at 2- or 3-day intervals.

### Nanoparticle dissolution study

Culture media from cellularized nanocomposite gel environments were collected and analyzed at day 3, 7, 14 and 21 to measure the released silicon concentration in solution. Samples of culture media from cellularized gels without nanoparticles were used as controls. Silicon content was measured by inductively-coupled plasma mass spectrometry (ICP-MS) using an Agilent 7900 ICPMS equipment.

### Cell viability, distribution and metabolic activity

Cell viability and distribution within the dense collagen gels were analysed by means of a Leica SP5 upright confocal (Leica DMI6000 Upright TCS SP5) coupled with a multiphoton laser scanning microscopy (Mai Tai multiphoton laser), which enabled the simultaneous acquisition of fluorescence and second-harmonic generation (SHG) signal. Prior to imaging, seeded cells were stained on day 28 post-culture following the live/dead® viability-cytotoxicity assay instructions (Thermofisher, Waltham, MA, USA). Constructs were washed in phosphate buffered saline (PBS) and stained with 2 µM calcein-AM and 4 µM ethidium homodimer-1 solution. Live cells were stained in green and dead cells in red. A hybrid collector with a wavelength of 438 ± 24 nm (initial excitation of 880 nm) was used to detect the SHG signal of the corresponding collagen fibers and a 25x/0.95 water objective with a working distance of 2.5 mm was used to acquire the z-stacks for each sample. Image series were acquired with the Leica Application Suite X software. Collagen signals were analysed using ImageJ.

Cell metabolic activity in dense collagen gels was analysed by alamarBlue® assay (Sigma-Aldrich, St. Louis, MO, USA) very week up to day 21 of culture. The cell-laden gels were incubated in a complete culture media containing 10 µg.mL^-1^ resazurin at 37°C/5% CO2 for 3 hours. Absorbances were read at wavelength of 570/600nm with a UV/Vis Uvikon spectrophotometer. The reduction percentage of AlamarBlue was calculated according to the manufacturer’s instructions.

### Scanning electron microscopy (SEM) / Energy-dispersive x-ray spectroscopy (EDS) studies

First, gels were fixed with 4% paraformaldehyde (PFA) and washed with 0.1M sodium cacodylate/0.6M sucrose buffer, before being dehydrated in a series of alcohol solutions and dried with supercritical CO2. Dried samples were mounted on metallic holders and sputter-coated with gold (15 nm layer) for SEM analysis and carbon (20 nm layer) for EDS. Samples were analysed under a Hitachi S-3400 N microscope at accelerating voltage of 10kV for SEM and 70 kV for EDS. Several pictures at different magnifications (1500-10,000x) were taken to examine microstructural features and chemical content.

### Histological and Immunohistochemistry staining

After 28 days in culture, gels were fixed in a 4% PFA solution, dehydrated, embedded in paraffin and cut into 7 µm thick sections. Prior to staining, sections were deparaffinized with toluene and rehydrated in ethanol solutions of decreasing concentrations, finishing with water. Cell distribution within the collagen matrix was assessed using Masson’s trichrome staining. Calcium and phosphate deposits were detected by alizarin red and Von Kossa, respectively. Micrographs were acquired under a light microscope (DMLB, Leica).

### qRT-PCR analyses

On days 1, 7, 14, and 21, cell-laden dense collagen gels were digested by collagenase, homogenized using Trizol® (Invitrogen, USA) and stored at -80°C prior to utilization. Total RNA content was extracted using the RNeasy Plus Mini Kit (Qiagen, Hilden, Germany) according to the manufacturer’s instructions. For quantitative PCR, single-stranded RNA was transformed into complementary double-stranded DNA (cDNA) by reverse transcription. 1 μL of random primers (200 uM) (Invitrogen) and 1 μL deoxyribonucleosides triphosphates (10 mM) (Invitrogen) were added to 10 µL RNA aliquots (around 300-500 ng). Following denaturation of secondary structure at 65°C and primers binding, 5X reaction buffer, dithiothreitol (0.2 M) and Moloney murine leukemia virus (M-MLV) reverse transcriptase (Invitrogen) were added. After 60 min at 37°C, the reaction was stopped by heating at 70°C for 10 min. The resulting cDNAs were stored at -20°C until further use.

Real-time fluorescence analysis using SYBR Green Master Mix (Roche Diagnostics GmbH, Germany) was used to quantify the expression level of specific genes relative to GAPDH or 18S rRNA housekeeping genes. The primers used in RT-qPCR are listed in **Supplementary Information, Table S1**. The reaction involved a succession of 40 cycles, each cycle including denaturation (95°C, 30 s), annealing (60°C, 30 s) and elongation with Taq polymerase (72°C, 30 s). Target gene quantities were normalized to GAPDH or18S rRNA using the Pfaffl method to determine variations in expression of each target gene between the different samples. Gels prepared without nanoparticles at day 7 were used as calibrator point. Three samples were analysed per group.

### Statistical analysis

Statistical analysis was performed on OriginPro 9.4 (OriginLab, Northampton, MA, USA). Data were compared statistically by one-way analysis of variance (ANOVA) tests. Statistical difference was considered at *p* < 0.05. Data are presented as mean ± standard deviation of mean (SD).

## RESULTS

### Nanoparticles and Nanocomposites Characterization

Spherical nanoparticles of silica (Si) and bioglass (BG), *ca*. 170 ± 50 nm and 140 ± 55 nm in diameter, were synthesized (**Figure 1A**). FTIR spectra of silica particles showed an intense band around 1100 cm^-1^ which corresponds to the stretching vibrations of the Si-O-Si bonds in the SiO4 tetrahedron, a band around 950 cm^-1^ corresponding to the Si-O bond of the silanols and a band around 800 cm^-1^ attributed to inter-tetrahedral Si-O-Si bending vibration mode (**Figure 1B**).^43^ For the bioglass particles, the band at 950 cm^-1^ almost disappeared, as a result of the heating treatment yielding to the condensation of the silicate network. WDXRF indicated that the bioglass particles displayed a Ca:Si atomic ratio of 0.05.

**Figure 1.**
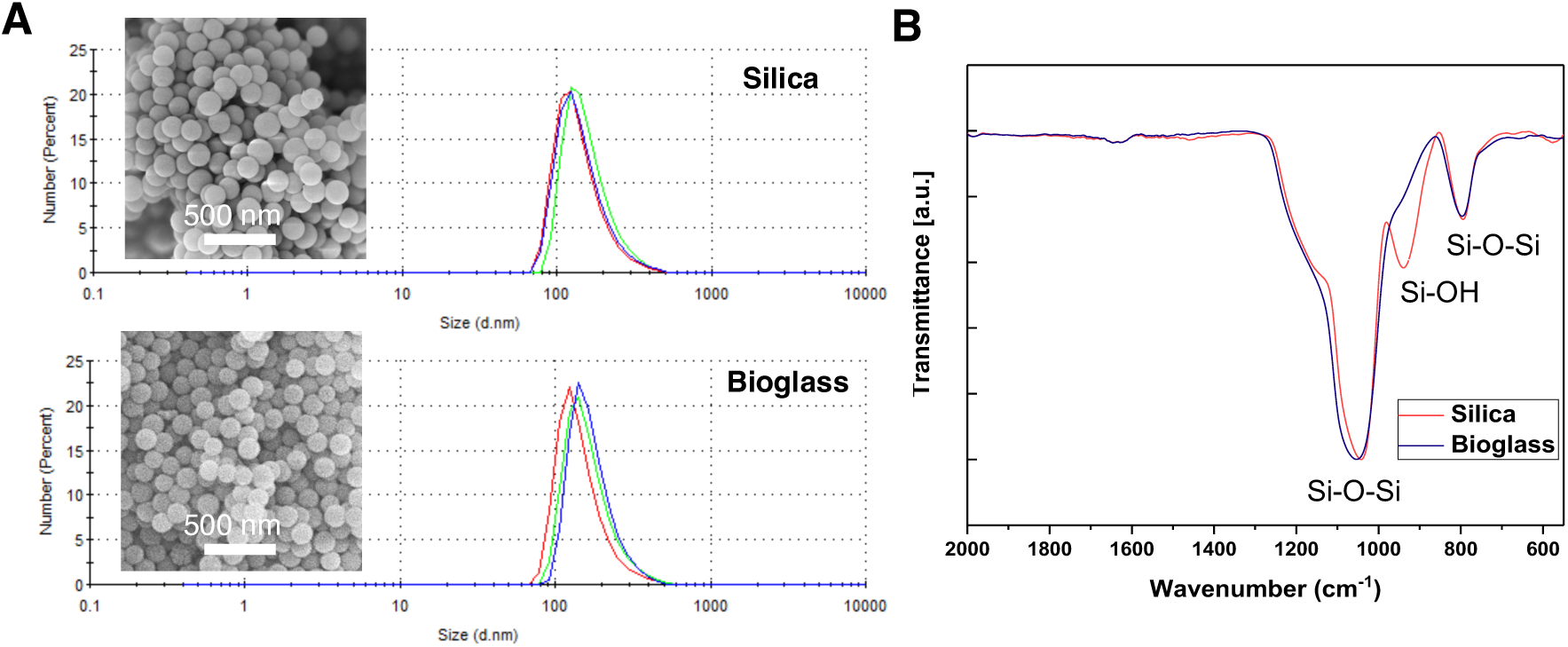
Characterization of the silica and bioglass particles. (A) DLS size distribution curves (3 measurements) and SEM images (inset, scale bar: 500 nm) and (B) FTIR spectra.

SEM images of as-prepared gels with and without particles at 100 μM and 1000 μM revealed a similar, dense, and random collagen fibril network, suggesting that nanoparticle presence does not disrupt fibril formation (**Figure 2A**). In parallel, ATR-FTIR spectra of as-made DC scaffolds exhibited characteristic collagen triple helix signals via amide I, II, and III bands at 1643, 1550, and 1243 cm^−1^, respectively (**Figure 2B**).^44^ No clear indication of Si-O-Si or PO4^3-^ groups in the 900-1100 cm^-1^ region was found on the spectra, likely due to their low concentrations in the gels.

**Figure 2.**
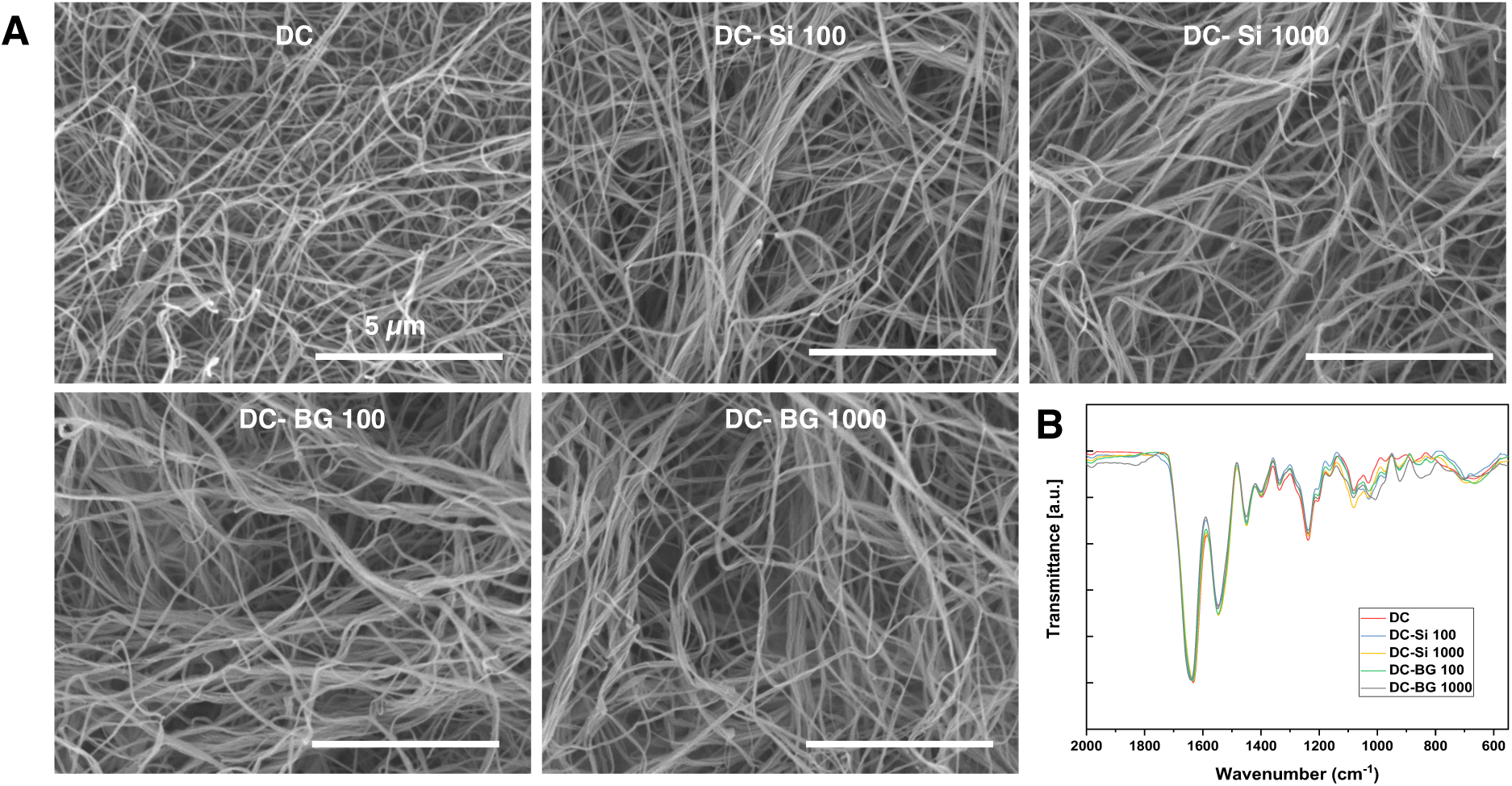
Morphological and chemical analyses of as-prepared acellular collagen gels (DC), nanocomposites with silica (Si) and bioglass (BG) nanoparticles at 100 μM and 1000 μM. (A) SEM images (scale bar: 5 μm) and (B) FTIR spectra.

The bioactivity of acellular gels was assessed after 21 days in mineralizing medium. with only DC-BG gels exhibiting mineral deposits along collagen fibrils (**Figure 3A**). EDS analysis confirmed that the deposited mineral particles contains both calcium and phosphorus with a Ca/P ratio of *ca*. 1.4, indicative of the formation of a poorly crystalline, substituted hydroxyapatite phase, as expected in the early stages of bioglass dissolution/reprecipitation (**Supplementary Information, Figure S1**).^45^ Moreover, the Si content was always below 0.5 wt % for both DC-Si and DC-BG samples, suggesting that only few, if any, nanoparticles remain. In parallel, FTIR spectra showed the presence of an intense asymmetric phosphate band between 900 and 1100 cm^−1^ for DC-BG (100 and 1000) gels, confirming mineralization of these samples while no peaks attributable to phosphate or silicate could be evidenced in the DC and DC-Si samples (**Figure 3B)**.

**Figure 3.**
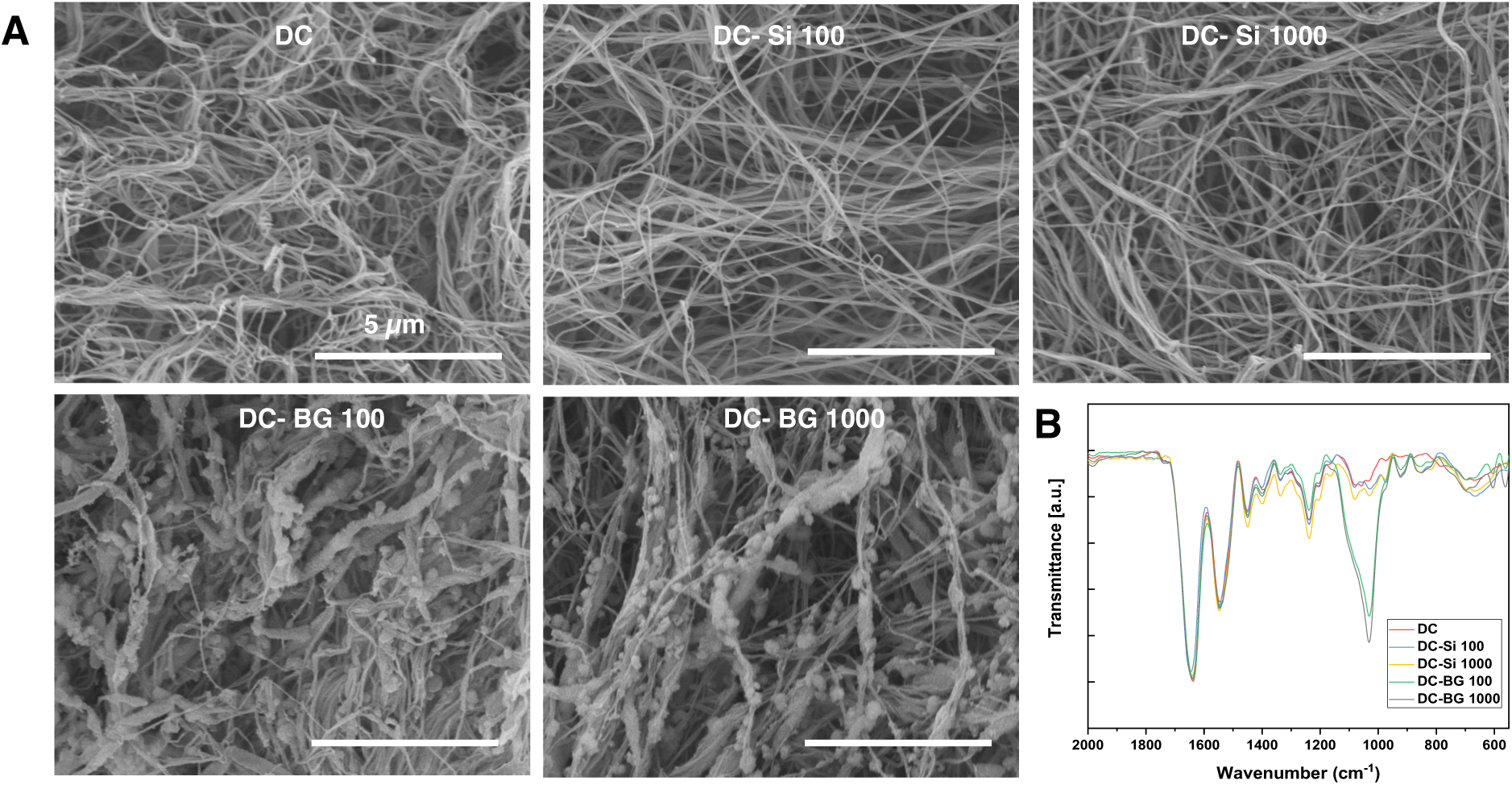
Morphological and chemical analyses of acellular collagen and nanocomposite hydrogels kept 21 days in mineralizing medium. (A) SEM images (scale bar: 5 μm) and (B) FTIR spectra.

### Influence of nanoparticles on hDPSC viability, metabolic activity and differentiation

The hDPSC-cellularized dense collagen DC hydrogels containing silica or bioglass nanoparticles at the same silicon concentrations, 100 μM and 1000 μM, were prepared and cultured up to 28 days in mineralizing induction medium (MIM). Live/Dead staining of hDPSCs was performed after incubation of 28 days. As shown in **Figure 4**, most hDPSCs were viable in the absence and presence of low (100 μM) and high (1000 μM) concentrations of silica or bioglass nanoparticles.

**Figure 4.**
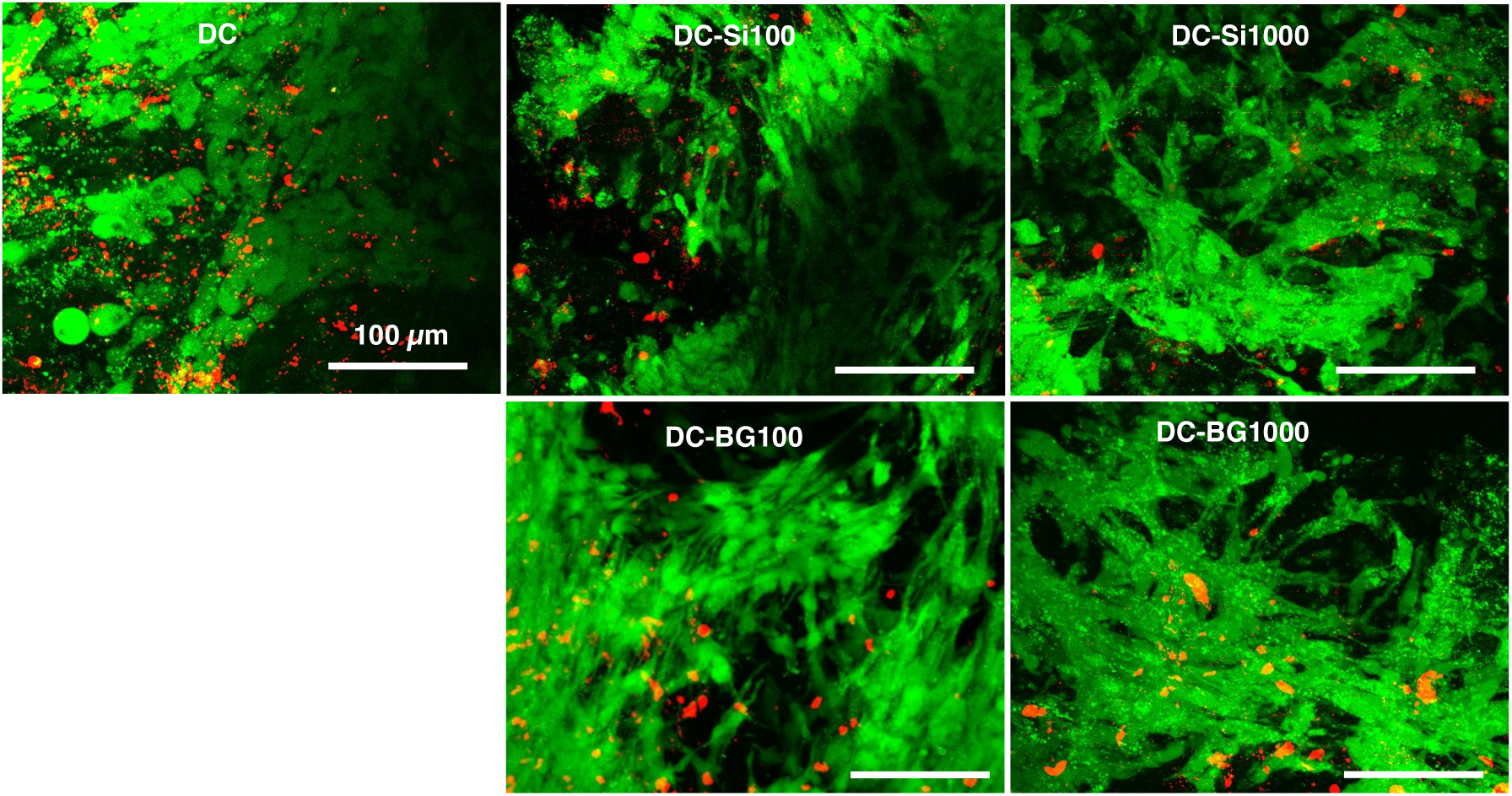
Confocal images of live (green) and dead (red) hDPSCs in collagen and nanocomposite hydrogels after 28 days of culture in mineralizing medium. (Scale bar: 100 μm).

Alamar Blue assays indicated that the metabolic activity of hDPSCs seeded in DC, DC-Si and DC-BG slowly decreased with culture time (**Figure 5A**). However, cells in DC-BG hydrogels displayed significantly higher metabolic activity than cells in DC alone and DC-Si at all time points. In parallel, the release of silicon in the gel supernatant, indicative of particle dissolution, was monitored by ICP-MS (**Figure 5B**). The release was the highest on day 3 in all the conditions, with mean released of concentrations of 100, 40, 20 and 10 μM in culture medium for DC-Si 1000, DC-BG 1000, DC-Si 100 and DC-BG 100, respectively. Dissolution extent remained larger for silica compared to bioglass nanoparticles at both concentrations until all curves levelled off at 10 μM, corresponding to the detection limit of the technique, after 3 weeks.

**Figure 5.**
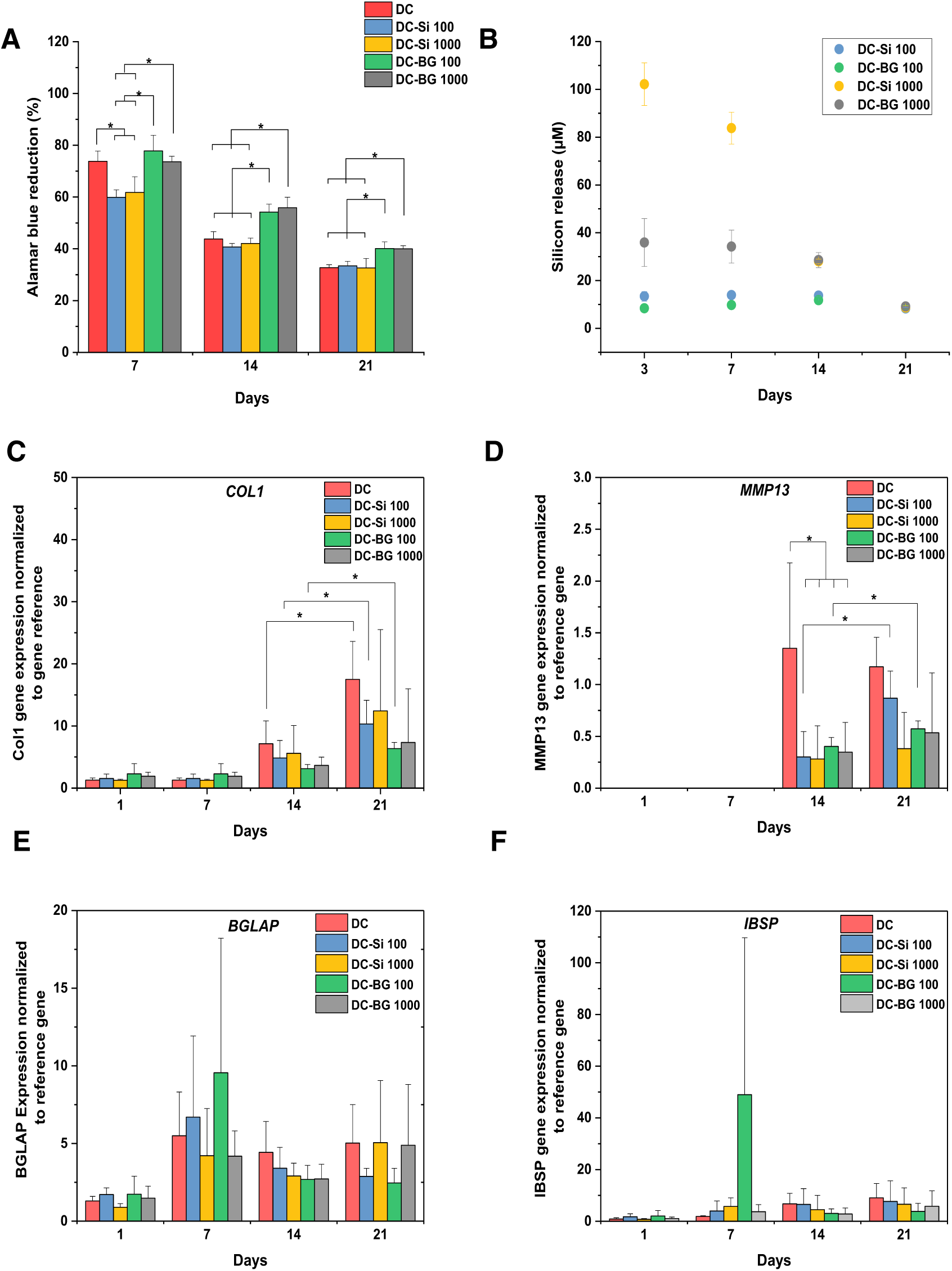
Metabolic activity, silicon release and cell differentiation over 21 days of culture in mineralizing medium of collagen and nanocomposite hydrogels. (A) hDPSC metabolic activity, as obtained from Alamar Blue assay, (B) silicon content in the supernatant, as measured by ICP-MS, and gene expression of (C) *COL1A1*, (D) *MMP13*, (E) *BGLAP* and (F) *IBSP* by qRT-PCR analysis. * p < 0.05

The expression of key genes involved in matrix colonization and remodeling, *COLI1A1* and *MMP13*, and mineralization, *BGLAP* (osteocalcin) and *IBSP*, was also followed over the whole culture period by qRT-PCR (**Figure 5(C-F)**). Expression of *BGLAP* and *IBSP* occurred at all times points and were not significantly different among different conditions. *COLI1A1* expression was low over the first week of culture and then increased with time for all samples, with no significant difference between the samples. *MMP13* expression was detected from day 14 and was lower in the presence of nanoparticles.

### Influence of nanoparticles on cell distribution and mineralization

Cell distribution within the collagen scaffolds was evaluated through Masson’s trichrome histological staining of gels cross-sections and SHG imaging coupled with confocal live-cell microscopy on day 28 of culture (**Figure 6**). Masson’s trichrome staining revealed distinct patterns of cell distribution within the gels. In cellularized DC and DC-BG 100 gels, clusters of cells were observed both inside and outside the gel, while they were predominantly located at the edges of DC-Si 100 and DC-Si 1000 gels (**Figure 6A)**. Conversely, a more uniform distribution of cells was observed throughout the DC-BG 1000 gel. The SHG/confocal 3D images further confirmed this uniform distribution of cells within the nano-composite gel loaded with 1000 µM bioglasses nanoparticles whereas large cell aggregates were observed in other samples (**Figure 6B)**. In parallel, SHG images of the collagen network were similar for all conditions, showing a globular organization with few large fibers (**Figure 6C)**. Qualitative analysis of fiber orientation showed limited variations between the samples (**Supplementary Information**, **Figure S2**).

**Figure 6.**
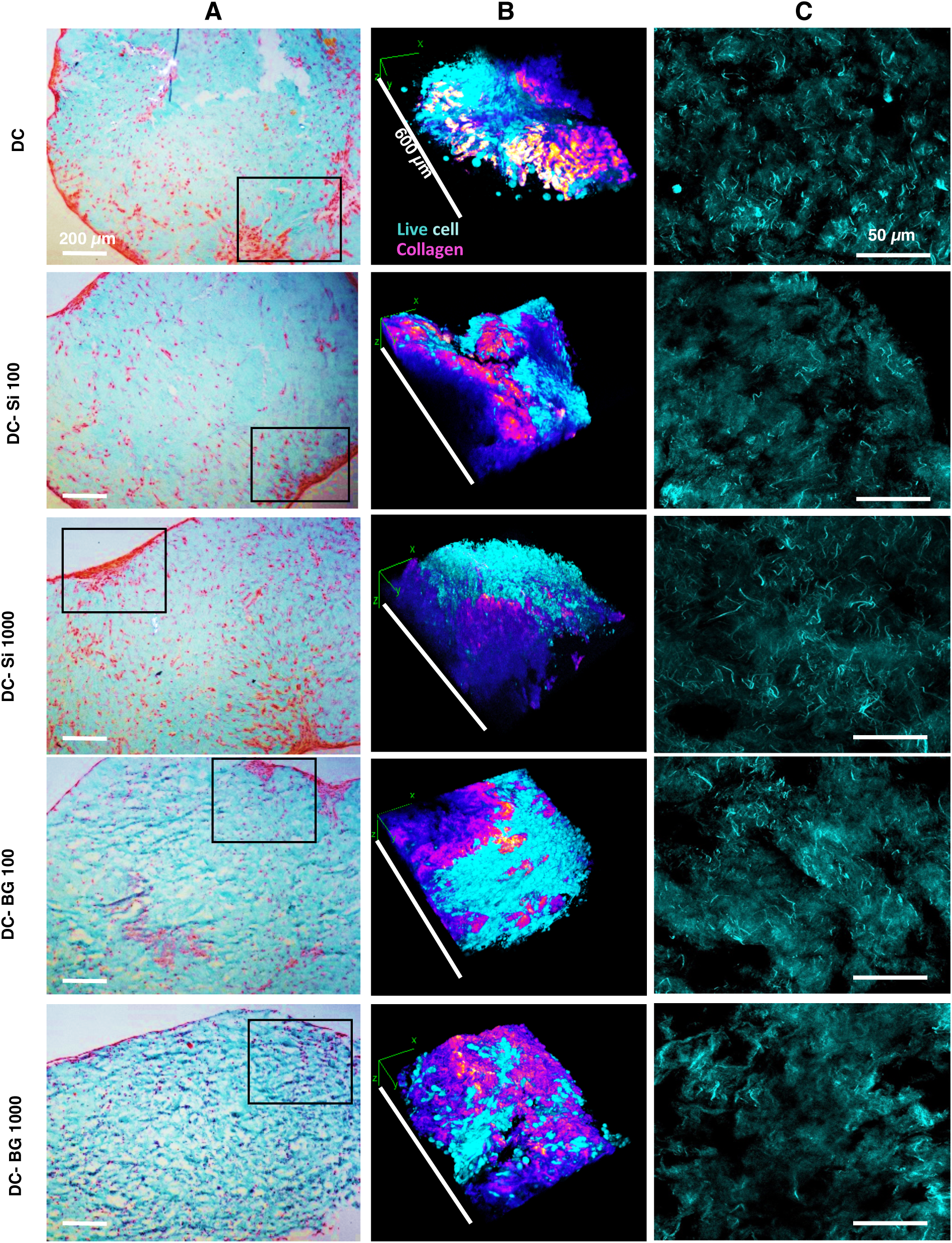
Cells distribution and matrix structure in collagen and nanocomposite hydrogels on day 28 of culture in mineralizing medium. (A) Masson’s Trichrome histological staining of hydrogel cross-sections (scale bar: 200 μm), (B) 3D rendering of confocal image coupled with SHG image for insets in (A) (collagen=purple, live cells=magenta; dead cells=red) (scale bar: 600 μm) and (C) SHG images of collagen (scale bar: 50 μm).

Von Kossa staining was used to detect calcium deposits in the gel sections, evidenced by dark coloration. All five types of cell-laden gel, with and without nanoparticles, were positive to Von Kossa staining after 28 days of culture in mineralizing medium, indicating the presence of minerals. However, the distribution of mineral deposits varied between gels. Like the distribution of cells revealed by Masson’s trichrome staining, mineral deposits occurred in clusters, corresponding to areas of cell aggregation, particularly evident with clustered minerals within DC gels and at the periphery of DC-Si 100 and DC-Si 1000 gels (**Figure 7**). Interestingly, the mineral deposit was slightly more uniform in the presence of 100 µM bioglasses and even more uniform in the presence of 1000 µM.

**Figure 7.**
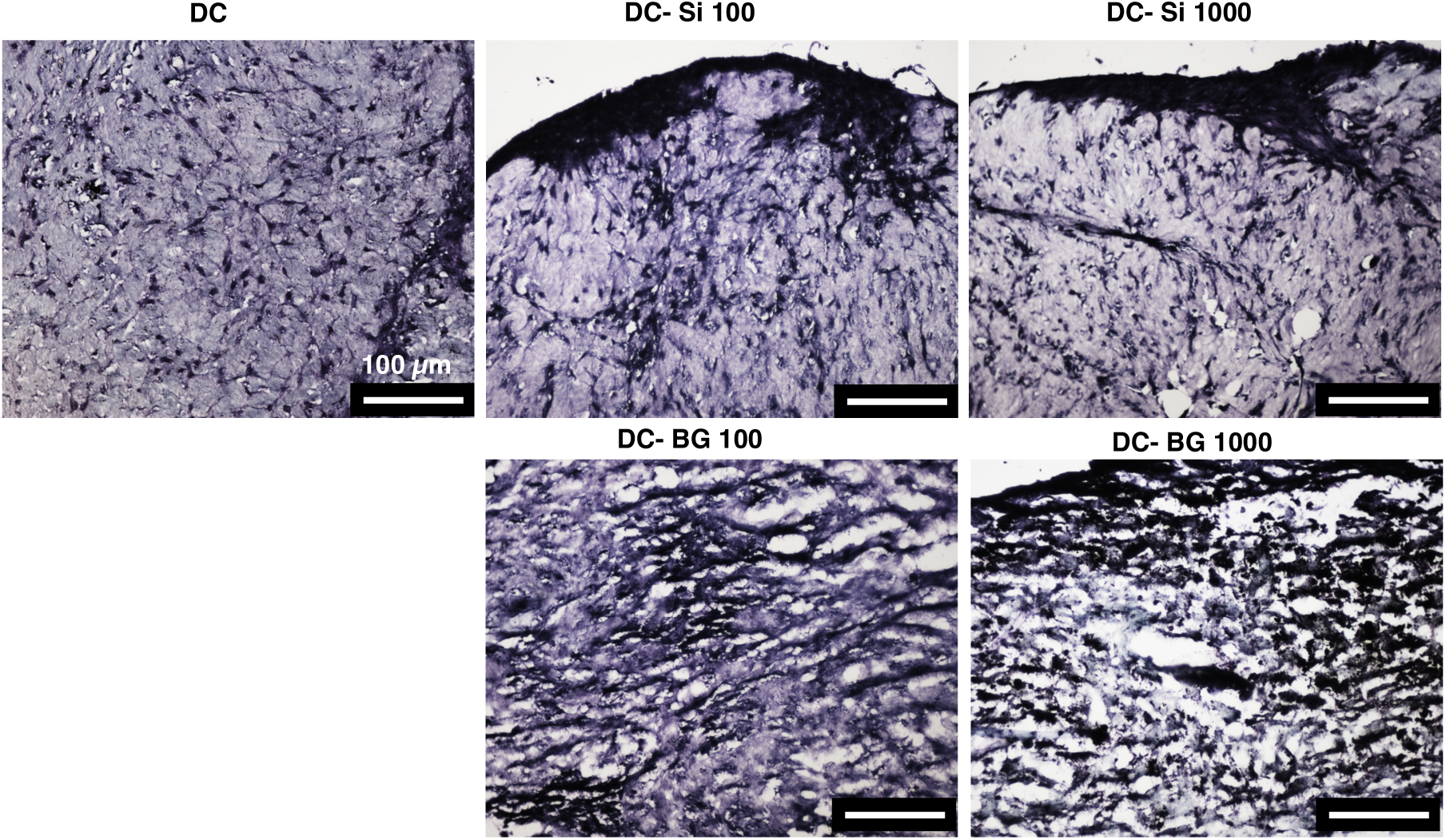
Von Kossa histological staining of cross-sections of collagen and nanocomposite hydrogels on day 28 of culture in mineralizing medium (scale bar : 100 μm).

SEM images of cellularized gels on day 28 also showed matrix mineralization across all gel types, with mineral deposition observed in front of cells and along collagen fibrils, suggesting cell-mediated mineralization **(Figure 8A**). In parallel, the FTIR spectra of all cell-seeded gels, regardless of the presence of nanoparticles (silica or bioglass) in the gels, exhibited a broad phosphate band between 900 and 1100 cm^−1^ (**Figure 8B**). No noticeable difference in mineralization extent was observed among the five groups, as phosphate band intensity was the similar in all samples.

**Figure 8.**
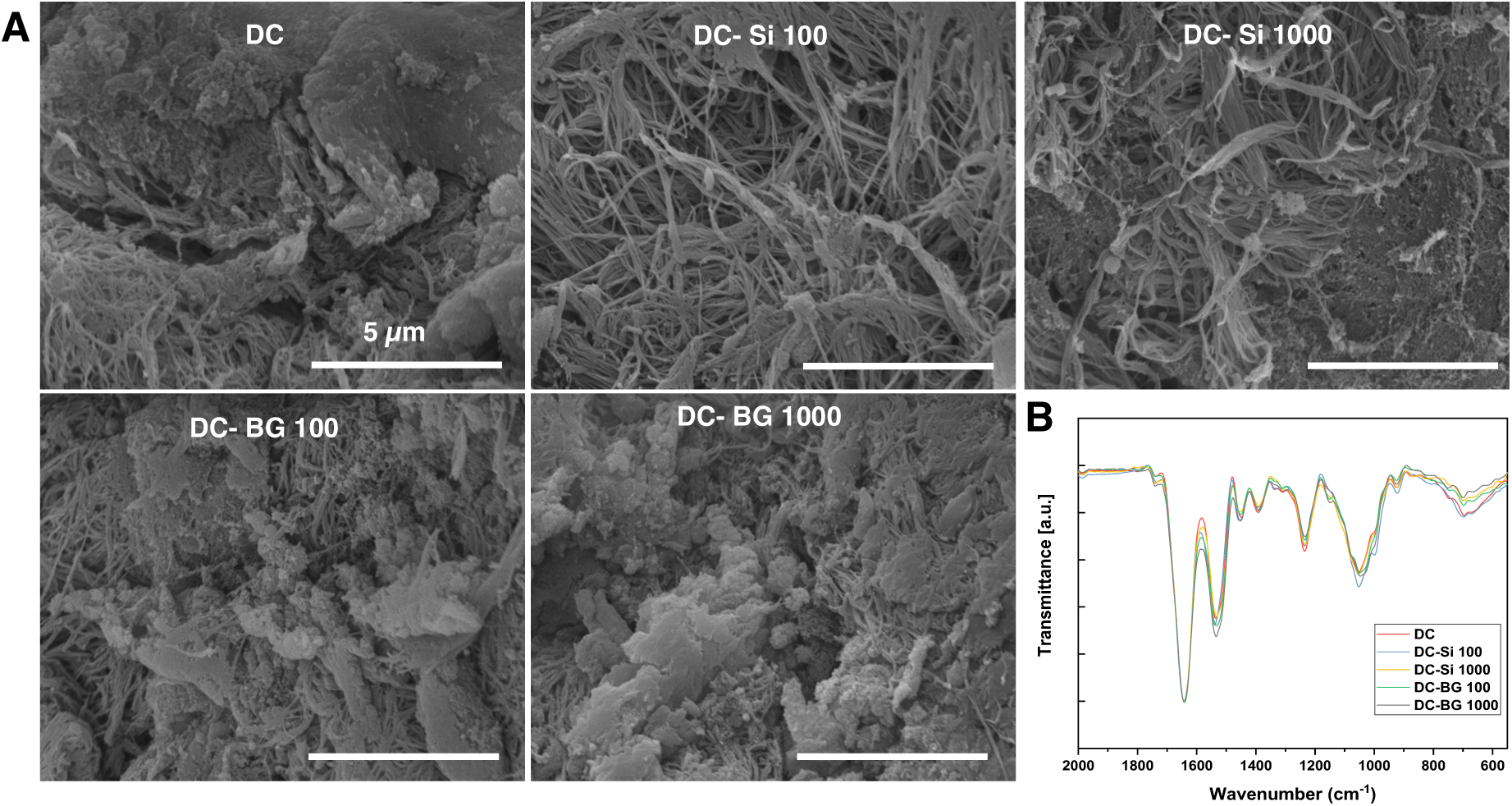
Mineralization of collagen and nanocomposite hydrogels on day 28 of culture in mineralizing medium. (A) SEM images (scale bar: 5 μm) and (B) FTIR spectra.

## DISCUSSION

In dentistry, silicon-based materials have large applications in dentin repair,^26^ although the specific impact of silicon on cells of the dentin-pulp complex is still poorly understood.^36–39^ It has been recently reported that the continuous addition of silicon could influence the viability and mineralization of human dental pulp stem cells (hDPSCs) within compressed 3D dense collagen hydrogels.^39^ However, in the clinical situation, silicon is progressively released as the result of the dissolution of the applied biomaterial, in contact with the dental tissue. To mimic this situation, silicon-releasing particles were here incorporated within the collagen hydrogels. Two types of nanoparticles, silica and bioglass, were studied in order to compare the effect of silicic acid release alone with the simultaneous release of both Si(OH)4 and Ca^2+^. Concentrations of 100 µM and 1000 µM in the collagen gels were selected as it was expected that their dissolution would lead to silicic acid concentration in the physiological-to-supraphysiological range, but below cytotoxic values.^36^

The sol-gel method allowed for the preparation of silica (Si) and bioglass (BG) nanoparticles of comparable size (*ca*. 150 nm in diameter). Their incorporation within dense collagen network at both 100 μm and 1 000 μm Si concentration did not result in modification of its fibrous structure, at least at the SEM scale, in agreement with the literature.^46^ The amount of calcium incorporated in the BG structure, although being low (Ca:Si < 0.1), was able to confer bioactive properties to the collagen hydrogel, whereas silica nanoparticles did not. Both types of particles released silicic acid from the nanocomposite gels over 3 weeks, the highest concentrations at a given time point being obtained for Si. This is due to the fact that BG preparation involves a high-temperature treatment leading to enhanced condensation of the silica network, thus decreasing its solubility rate. Noticeably, released silicic acid concentration from DC-BG1000 and DC-Si1000 was above the reported range physiological values over the first two weeks.

While no clear effect of the particles on the cell survival could be evidenced by Live/Dead staining, their metabolic activity after 1 week of culture was decreased by Si particles but increased by BG particles. The fact that such variations do not depend on the released Si(OH)4 concentrations (10-100 μM), that were previously shown not to decrease DPSCs metabolic activity, suggests that cell-particles interactions are to be considered. Silica NPs can be taken up by cells and, depending on their size, influence cellular functions.^47^ For example, it was shown that 80 nm SiO2 nanoparticles internalized by cells induced a significant loss of mitochondrial membrane integrity and a decrease in mitochondrial dehydrogenase activity, which could directly impact cellular metabolism.^48^ In contrast, internalized BG particles could be favorable to bone marrow mesenchymal stem cells activity.^49^

The differences observed in cell distribution within the collagen matrix provide valuable insights into the impact of Si and BG on cellular behavior. First, cells clusters appeared both inside and outside DC gels while cells were mainly located at the edges of DC-Si 100 and DC-Si1000 gels. This suggests that silica nanoparticles can influence cell migration and distribution within the collagen scaffolds, consistent with previous observations showing that nanoparticles uptake by fibroblasts could impair their adhesion and migration.^48^ A second possible explanation is that released silicic acid interacts with the collagen molecules.^50^ This interaction could affect local cell adhesion to the collagen network, favoring hDPSC migration to the external parts of the gels. It is important to point out that, while cell clustering was previously observed in the presence of silicic acid only, the highly heterogeneous distribution of hDPSCs seems specific to the presence of silica in a particulate form. In contrast, BG NPs led to a more homogeneous cell dispersion over the whole gel volume, in a concentration-dependent manner. This would suggest that the release of calcium can favour local hDPSCs proliferation, this effect prevailing over the silicon influence. Importantly, a delay in the expression of the MMP13 gene was observed in all nanocomposite gels, in line with the previous findings using constant silicic acid supplementation.^39^ MMP13 is a key protein in matrix remodelling, a process that requires cell adhesion and migration.^51^ Hence its lower expression level would support the hypothesis that particles and/or released silicic acid affect cell-matrix interactions.

Regarding mineral deposition, all gel groups were mineralized by the end of the culture time in the presence of hDPSCs and the distribution of mineral deposits matched/mirrored the observed distribution of cells in the gels. Moreover, the gene expression pattern of key mineralizing proteins was similar in all conditions and FTIR data suggest that the extent of mineralization was similar in all groups. Taken together, these data suggest that mineralization proceeds by a cell-mediated process in hDPSC-seeded collagen gels, with no significant influence of silicic acid. This strengthens the hypothesis that Si(OH)4 has no direct effect of the mineralizing ability of hDPSCs in the 10-100 μM range.^39^ Furthermore the cell-mediated mineralization did not appeared influenced by the intrinsic bioactivity of the BG nanoparticles, which may be related to their low calcium content.^49^

ATR-FTIR analysis revealed a more intense phosphate band in acellular than in cellularized DC-BG gels. Although such comparison must be taken cautiously as the shape of the bands was not the same in both series of samples, it would suggest that, in the conditions of this work, cell-free constructs may outperform cell-seeded ones in mineralization. Similar findings were previously observed *in vivo* where an acellular DC-BG collagen gel implanted in critical-sized murine calvaria defect resulted in a greater bone volume than a DPSC-cellularized DC-BG collagen gel.^52^ This suggests that, *in vivo*, the chemically-induced hydroxyapatite deposition from bioglass may be particularly efficient in recruiting host mineralizing cells while cell-induced mineralization inside the scaffold may be less favorable for further interaction with the surrounding tissue. The cross-talk between implanted and host cells can also contribute to this difference.

Altogether, these results confirm that silicic acid at concentrations below 100 μM has no significant influence on DPSCs survival or mineralization ability in 3D dense collagen hydrogels. Moreover, they highlight the fact that the presence of silica nanoparticles and their dissolution product, Si(OH)4, have a marked influence on cell adhesion, migration and matrix remodelling ability. This may result from a combination of biological interactions of the particles with cells and chemical interactions of soluble silica with collagen. Conversely, the fact that the bioglass do differs in its influence could be attributed to the absence of reported effects of BG internalization on cell adhesion and migration as well as to the possibility that calcium ions may prevent Si(OH)4-collagen interactions. Such an ability of Ca^2+^ to regulate the reactivity of silicic acid could contribute to the current gap existing between *in vivo* experiments where the benefits of silicic acid release in bone repair has been well-established and *in vitro* studies that are overall less conclusive.^36^

## CONCLUSION

In conclusion, this study demonstrates that the effect of silicon-releasing nanomaterials on the *in vitro* mineralization of pulp-like living materials results from a subtle interplay between chemical and biological processes, the former being often underrated in the literature. While this conclusion could apply to many other biomaterials and tissues, the context of this study has several specificities: (i) there is, to date, no proof of the possible role of silicon in dental tissue formation or physiological repair, (ii) while the interaction of silica particles with cells and biomolecules has been extensively studied, the mechanisms by which silicic acid can interact with living tissues at the molecular level remain poorly understood. Therefore, although using currently-applied biomaterials and/or designing pulp models of increased biomimetic character may enhance the relevance of such *in vitro* studies to clinical situations, it is also important to pursue more fundamental approaches using simple models.

## ASSOCIATED CONTENT

### Supporting Information

The following files are available free of charge.

Primers for PCR; SEM/EDX analysis of acellular samples; quantitative analysis of collagen fiber orientation (PDF)

## AUTHOR INFORMATION

### Author Contributions

The manuscript was written through contributions of all authors. All authors have given approval to the final version of the manuscript.

### Funding Sources

This work was funded by the French Ministry of Superior Education and Research (Doctorate School 397). This source had no involvement in work conduction and manuscript submission

## ACKNOWLEDGMENT

The authors thank I. Génois (LCMCP) for WXRDF analyses and P. Burkel (Institut de Physique du Globe de Paris) for ICP-MS analyses. T.C. thanks Dr A.-M. Collignon (UR2496, Université Paris Cité) for fruitful discussions.

## Notes

### Competing Interest Statement

The authors have declared no competing interest.

