## Supporting Information for "Evaluation of Silica and Bioglass Nanomaterials in Pulp-Like Living Materials"

**Table S1**. Primers for real-time PCR

**Figure S1**. SEM/EDX of acellular samples

**Figure S2.** Quantitative analysis of collagen fiber orientation

**Table S1**. Primers for real-time PCR

| Primer | Sequence (Forward 5'- 3') | Sequence (Reverse 5'- 3') | Supplier |
|---|---|---|---|
| BGLAP | CAAAGGTGCAGCCTTTGTGTC | TCACAGTCCGGATTGAGCTCA | Roche Diagnostics GmbH, Germany |
| Col1A1 | CCAGTCAGAGTGGCACATCTTGA | GCTCACGATGGTGCCGCTACTA | |
| MMP13 | GCCGGTGTAGGTGTAGATAGGAAA | GGAGATGCCCATTTTGATGATGA | |
| IBSP | AACGAACAAGGCATAAACGGCACCA | CTTGCCCTGCCTTCCGGTCT | |
| ALP | GCAGCTTGACCTCCTCGGAAGACACT | TCACCGCCCACCACCTTGTAGCC | |
| GAPDH | TTGATTTTGGAGGGATCTCG | GAGTCAACGGATTTGGTCGT | |
| 18S rRNA | TTACAGGGCCTCGAAAGAGT | TGAGAAACGGCTACCACATC | |

**Figure S1**. SEM/EDX analysis of acellular hydrogels after 21 days in mineralizing ledium

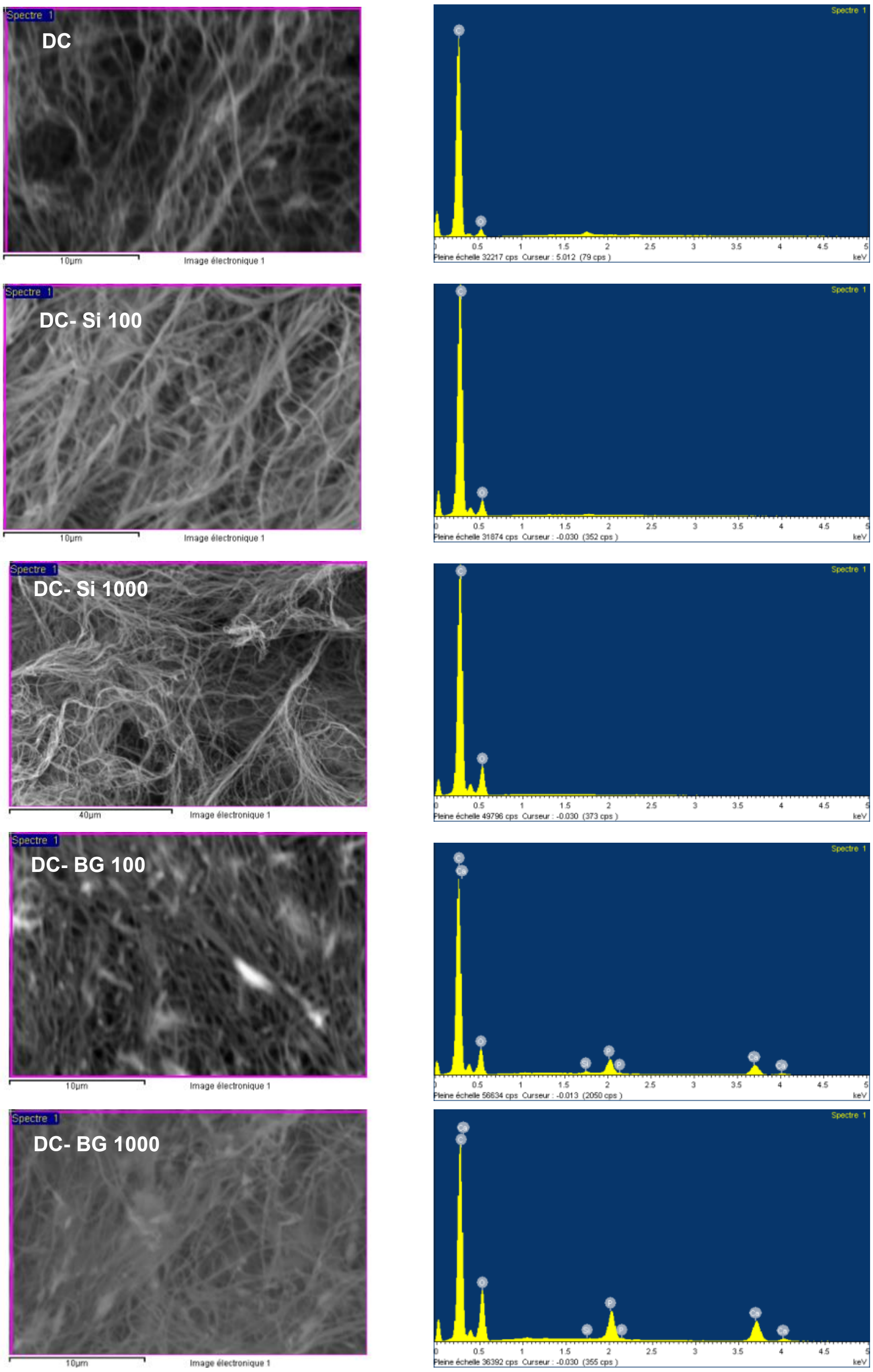

**Figure S2**. Quantitative analysis of collagen fiber orientation from SHG images of DC, DC-Si and DC-BG hydrogels on day 28 of culture.

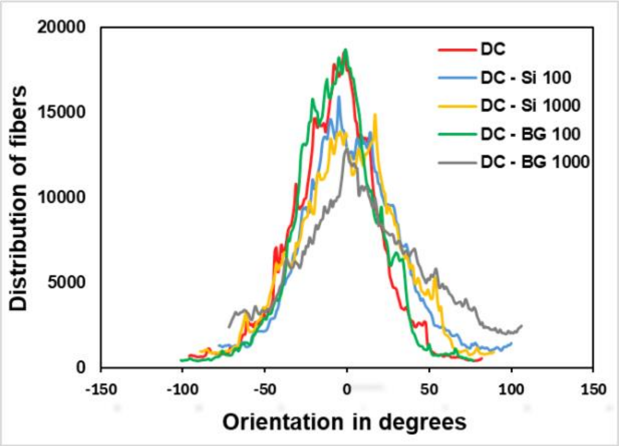
